## Supplementary Tables for "Egg-adaptation pathway of human influenza H3N2 virus is contingent on natural evolution"

**Supplementary Table 1. H3N2 components of WHO-recommended influenza vaccines for influenza seasons between 2008 and 2020.**

| SEASON | STRAIN NAME | REFERENCE |
|---|---|---|
| 2008-2010 | A/Brisbane/10/2007 | https://www.who.int/influenza/vaccines/200809Recommendation.pdf |
| 2010-2011 | A/Perth/16/2009 (NIB-64) | https://www.who.int/influenza/vaccines/virus/recommendations/201002_Recommendation.pdf |
| 2011-2012 | | https://www.who.int/influenza/vaccines/2011_02_recommendation.pdf |
| 2012-2013 | A/Victoria/361/2011 (X-217, IVR-165) | https://www.who.int/influenza/vaccines/virus/recommendations/201202_recommendation.pdf |
| 2013-2014 | | https://www.who.int/influenza/vaccines/virus/recommendations/201302_recommendation.pdf |
| 2014-2015 | A/Texas/50/2012 (X-223) | https://www.who.int/influenza/vaccines/virus/recommendations/201402_recommendation.pdf |
| 2015-2016 | A/Switzerland/9715293/2013 (NIB-88, X-247, IVR-176) | https://www.who.int/influenza/vaccines/virus/recommendations/201502_recommendation.pdf |
| 2016-2017 | A/Hong Kong/4801/2014 (X-263) | https://www.who.int/influenza/vaccines/virus/recommendations/201602_recommendation.pdf |
| 2017-2018 | | https://www.who.int/influenza/vaccines/virus/recommendations/201703_recommendation.pdf |
| 2018-2019 | A/Singapore/INFIMH-16-0019/2016 (IVR-186, NYMC-X-307, NIB-104) | https://www.who.int/influenza/vaccines/virus/recommendations/201802_recommendation.pdf |
| 2019-2020 (Northern hemisphere) | A/Kansas/14/2017 (X-327, IVR-195) | https://cdn.who.int/media/docs/default-source/influenza/cvvs/cvv-northern-hemisphere-2019-2020/summary-a-h3n2-cvv-egg-nh1920.pdf |
| 2019 (Southern hemisphere) | A/Switzerland/8060/2017 (NIB-112) | https://www.who.int/influenza/vaccines/virus/recommendations/201809_recommendation.pdf |
| 2020-2021 | A/Hong Kong/2671/2019 (NIB-121) | https://www.who.int/influenza/vaccines/virus/recommendations/202002_recommendation.pdf |

**Supplementary Table 2. H3N2 genes that were used in this study.**

| PARENTAL VIRUS | STRAIN NAME | CLADE | GENE | EGG-ADAPTIVE MUTATION | GISAID ACCESSION NO. |
|---|---|---|---|---|---|
| A/Kansas/14/2017 | X-327 | 3C.3a1 | HA | G186V, D190N, S219Y | EPI1490170 |
| | | | NA | - | EPI1504534 |
| A/Switzerland/8060/2017 | NIB-112 | 3C.2a2 | HA | T160K, L194P | EPI1313524 |
| | | | NA | - | EPI1313525 |
| A/Singapore/INFIMH-16-0019/2016 | WT | 3C.2a1 | HA | - | EPI1381186 |
| | | | NA | - | EPI1381185 |
| A/Italy/11871/2020 | WT | 3C.2a1b.1a | HA | - | EPI1735610 |
| | | | NA | - | EPI1735609 |
| A/Victoria/22/2020 | WT | 3C.2a1b.1a | HA | - | EPI1721444 |
| | | | NA | - | EPI1721443 |

‘-‘: no egg-adaptive mutation.

**Supplementary Table 3. Frequencies of mutations in egg-passaged viruses that showed an average frequency of >10% in the fifth passage.**

| INOCULUM | MUTATION | BIOLOGICAL REPLICATE | FREQUENCY DURING EGG-PASSAGING | | | | |
|---|---|---|---|---|---|---|---|
| | | | E1 | E2 | E3 | E4 | E5 |
| Kansas17 V186/L194/N190/Y219 (X-327) | - | Rep1 | - | - | - | - | - |
| | | Rep2 | - | - | - | - | - |
| | | Rep3 | - | - | - | - | - |
| Switz17 K160/G186/P194 | T203I | Rep1 | 0.01 | 0.62 | 0.97 | 0.98 | 0.98 |
| | | Rep2 | 0.06 | 0.14 | 0.32 | 0.53 | 0.85 |
| | | Rep3 | 0.08 | 0.13 | 0.23 | 0.57 | 0.75 |
| Sing16 K160/G186/P194 | I140K | Rep1 | 0.00 | 0.00 | 0.00 | 0.00 | 0.00 |
| | | Rep2 | 0.00 | 0.03 | 0.49 | 0.48 | 0.73 |
| | | Rep3 | 0.00 | 0.00 | 0.00 | 0.00 | 0.00 |
| | T203I | Rep1 | 0.01 | 0.01 | 0.07 | 0.21 | 0.27 |
| | | Rep2 | 0.45 | 0.16 | 0.43 | 0.49 | 0.24 |
| | | Rep3 | 0.01 | 0.01 | 0.02 | 0.57 | 0.90 |
| Kansas17 X-327 V186G/L194P/N190D | I140K | Rep1 | 0.00 | 0.00 | 0.00 | 0.02 | 0.06 |
| | | Rep2 | 0.15 | 0.44 | 0.71 | 0.82 | 0.87 |
| | | Rep3 | 0.02 | 0.05 | 0.14 | 0.19 | 0.28 |
| | T203I | Rep1 | 0.37 | 0.70 | 0.88 | 0.86 | 0.93 |
| | | Rep2 | 0.00 | 0.01 | 0.01 | 0.01 | 0.00 |
| | | Rep3 | 0.03 | 0.07 | 0.17 | 0.22 | 0.21 |
| | D225N | Rep1 | 0.00 | 0.00 | 0.02 | 0.10 | 0.28 |
| | | Rep2 | 0.01 | 0.02 | 0.02 | 0.01 | 0.01 |
| | | Rep3 | 0.02 | 0.10 | 0.25 | 0.34 | 0.39 |

Continued

| INOCULUM | MUTATION | BIOLOGICAL REPLICATE | FREQUENCY DURING EGG-PASSAGING | | | | |
|---|---|---|---|---|---|---|---|
| | | | E1 | E2 | E3 | E4 | E5 |
| Switz17 K160/V186/L194 | - | Rep1 | - | N/A | N/A | N/A | N/A |
| | | Rep2 | - | N/A | N/A | N/A | N/A |
| | | Rep3 | - | N/A | N/A | N/A | N/A |
| Sing16 K160/V186/L194 | H156R | Rep1 | 0.00 | 0.00 | N/A | N/A | N/A |
| | | Rep2 | 0.00 | 0.98 | N/A | N/A | N/A |
| | | Rep3 | 0.00 | 0.02 | N/A | N/A | N/A |
| | D190N | Rep1 | 0.00 | 0.00 | N/A | N/A | N/A |
| | | Rep2 | 0.00 | 0.00 | N/A | N/A | N/A |
| | | Rep3 | 0.00 | 0.94 | N/A | N/A | N/A |
| | D225N | Rep1 | 0.12 | 0.45 | N/A | N/A | N/A |
| | | Rep2 | 0.00 | 0.00 | N/A | N/A | N/A |
| | | Rep3 | 0.40 | 0.98 | N/A | N/A | N/A |

'-': no mutation found. N/A: copy number of viral RNA was insufficient for deep sequencing analysis.

**Supplementary Table 4. Information of HA from human H3N2 strains with or without egg-passaging.**

| VIRUS NAME | NON-EGG-PASSAGED | | EGG-PASSAGED | | MUTATION IN EGG-PASSAGED | | | | | |
|---|---|---|---|---|---|---|---|---|---|---|
| | PASSAGE HISTORY | HA ACCESSION NO. | PASSAGE HISTORY | HA ACCESSION NO. | 160 | 186 | 190 | 194 | 219 | 225 |
| A/Bangladesh/3005/2020 | Original | EPI1838357 | E3 | EPI1844062 | I | N | N | L | S | G |
| | | | E3+E6 | EPI1857491 | I | N | N | L | S | G |
| A/Bangladesh/3011/2020 | Original | EPI1838349 | E3 | EPI1844060 | I | N | N | L | S | G |
| A/Bangladesh/4002/2020 | Original | EPI1838416 | E3 | EPI1844058 | I | N | N | L | S | G |
| A/Bangladesh/911009/2020 | Original | EPI1838408 | E3+E6 | EPI1857489 | I | N | N | L | S | G |
| A/Beijing-Daxin/33/2020 | C2 | EPI1753671 | E7 | EPI1753663 | K | D | N | L | S | D |
| A/Beijing-Miyun/51/2020 | C1 | EPI1753655 | E7+E2 | EPI1805765 | A | D | N | L | Y | G |
| | | | E9 | EPI1801276 | I | D | N | L | F | G |
| A/Beijing-Miyun/53/2020 | C1 | EPI1753639 | E5+E2 | EPI1806767 | I | D | N | L | F | G |
| A/Beijing-Miyun/54/2020 | C1 | EPI1753623 | E4 | EPI1753615 | T | V | N | L | S | G |
| A/Belgium/G0023/2019 | SIAT1 | EPI1436935 | E3 | EPI1592043 | K | G | D | L | S | G |
| A/Brisbane/148/2019 | SIAT1+SIAT1 | EPI1658656 | E5 | EPI1740888 | K | V | D | L | Y | D |
| A/Brunei/39/2020 | SIAT1 | EPI1797893 | E5 | EPI1804363 | K | D | N | L | S | D |
| A/California/194/2019 | Original | EPI1630686 | E5 | EPI1713739 | K | V | D | L | S | D |
| A/Cambodia/e0826360/2020 | Original | EPI1837753 | E5+E1+E8 | EPI1877927 | K | R | D | L | F | D |
| A/Canberra/407/2019 | SIAT1 | EPI1671923 | E4 | EPI1713737 | K | G | D | P | S | D |
| A/Christchurch/515/2019 | SIAT2 | EPI1484393 | E3+E2 | .2a1b.2b | T | G | D | P | F | D |
| A/Christchurch/516/2019 | SIAT1 | EPI1484453 | E3 | EPI1491158 | K | V | D | L | F | D |
| A/Darwin/1/2021 | Original | EPI1851811 | E8 | EPI1859990 | K | S | D | L | S | G |
| A/Darwin/11/2021 | Original | EPI1859986 | E3 | EPI1859998 | I | D | N | L | S | D |
| A/Darwin/17/2021 | SIAT1 | EPI1888120 | E3 | EPI1924410 | I | V | N | L | S | D |
| A/Darwin/2/2021 | Original | EPI1851819 | E3 | EPI1859970 | K | R | D | L | S | D |
| A/Darwin/22/2021 | SIAT1 | EPI1888104 | E3 | EPI1923164 | I | N | N | L | S | G |

Continued

| VIRUS NAME | NON-EGG-PASSAGED | | EGG-PASSAGED | | MUTATION IN EGG-PASSAGED | | | | | |
|---|---|---|---|---|---|---|---|---|---|---|
| | PASSAGE HISTORY | HA ACCESSION NO. | PASSAGE HISTORY | HA ACCESSION NO. | 160 | 186 | 190 | 194 | 219 | 225 |
| A/Darwin/24/2021 | Original | EPI1888096 | E3 | EPI1923174 | I | N | N | L | S | G |
| A/Darwin/402/2019 | SIAT1 | EPI1508629 | E3 | EPI1584616 | T | G | D | P | F | N |
| A/Darwin/6/2021 | Original | EPI1857216 | E3+E7+E1 | EPI1885098 | I | N | N | L | S | G |
| | | | E5+E2 | EPI1925255 | I | D | N | L | F | G |
| A/Darwin/726/2019 | SIAT1 | EPI1658695 | E6 | EPI1675460 | A | V | E | L | S | D |
| A/Darwin/9/2021 | SIAT1 | EPI1883349 | E4 | EPI1888006 | I | N | N | L | S | G |
| A/Delaware/01/2021 | Original | EPI1869534 | E2 | EPI1940656 | I | D | N | L | S | G |
| A/Finland/183/2020 | SIAT1 | EPI1753460 | E6 | EPI1847912 | K | S | D | L | S | D |
| A/Hong Kong/2671/2019 | MDCK1 | EPI1543098 | E9 | EPI1843071 | I | V | D | L | F | N |
| A/Kansas/14/2017 | SIAT2 | EPI1504535 | E17 | EPI1415371 | K | V | N | L | Y | D |
| A/KANAGAWA/ZC1841/2019 | SIAT1 | EPI1478189 | E7 | EPI1696421 | K | V | D | L | S | D |
| A/Michigan/173/2020 | Original | EPI1843859 | E4+E2 | EPI1922037 | I | N | N | L | S | G |
| A/Netherlands/00007/2021 | Original | EPI1885138 | E3 | EPI1924781 | I | N | N | L | S | G |
| A/Newcastle/42/2019 | SIAT1 | EPI1430423 | E3 | EPI1444940 | K | V | D | L | F | D |
| A/Newcastle/623/2019 | SIAT1 | EPI1430438 | E2 | EPI1444941 | K | G | D | L | S | G |
| A/Norway/16606/2021 | SIAT1 | EPI1922181 | E3 | EPI1940648 | I | N | N | L | S | G |
| A/Norway/2279/2019 | SIAT1 | EPI1619464 | E4 | EPI1719268 | K | G | D | P | S | D |
| A/Oregon/28/2019 | Original | EPI1627961 | E6 | EPI1713733 | K | V | D | L | S | D |
| A/Paris/2554/2019 | Original | EPI1638885 | E4+E7 | EPI1794629 | I | D | N | L | S | G |
| A/Pennsylvania/01/2021 | Original | EPI1858654 | E5 | EPI1924787 | I | G | N | L | S | G |
| | | | E4 | EPI1924789 | I | D | N | L | S | G |

Continued

| VIRUS NAME | NON-EGG-PASSAGED | | EGG-PASSAGED | | MUTATION IN EGG-PASSAGED | | | | | |
|---|---|---|---|---|---|---|---|---|---|---|
| | PASSAGE HISTORY | HA ACCESSION NO. | PASSAGE HISTORY | HA ACCESSION NO. | 160 | 186 | 190 | 194 | 219 | 225 |
| A/Pennsylvania/1025/2019 | Original | EPI1630907 | E3+E1 | EPI1796140 | I | V | D | L | S | N |
| | | | E3+D8+E1 | EPI1804937 | I | V | E | L | S | D |
| | | | E3 | EPI1713729 | I | V | D | L | S | D |
| A/Pennsylvania/1026/2019 | Original | EPI1631152 | E5+E8 | EPI1794631 | K | V | D | L | Y | D |
| | | | E5+E2+E9 | EPI1843569 | K | V | D | L | Y | D |
| A/Perth/20/2020 | MDCK1+SIAT1 | EPI1733852 | E3+E2+E9 | EPI1848094 | K | D | N | L | Y | G |
| | | | E3 | EPI1740884 | K | D | N | L | Y | D |
| | | | E3+E7 | EPI1794633 | K | D | N | L | Y | N |
| A/Saitama/92/2020 | MDCK1+hMDCK1 | EPI1847848 | E4 | EPI1847862 | K | R | D | L | S | D |
| A/Singapore/INFKK0001/2021 | Original | EPI1883806 | E4 | EPI1889199 | I | N | N | L | S | G |
| A/Singapore/INFKK0002/2021 | Original | EPI1883814 | E4 | EPI1924397 | I | N | N | L | S | G |
| A/Singapore/KK0001/2020 | Original | EPI1750786 | E3 | EPI1804361 | K | D | D | L | S | D |
| A/South Africa/R06421/2019 | MDCK1+SIAT1 | EPI1582733 | E4 | EPI1694137 | K | V | D | L | F | D |
| A/South Australia/2/2019 | SIAT1 | EPI1387412 | E4+E2 | EPI1698481 | I | D | D | L | S | D |
| A/South Australia/320/2019 | SIAT1 | EPI1484436 | E4 | EPI1526498 | T | G | D | P | F | D |
| A/South Australia/34/2019 | SIAT1 | EPI1387331 | E5 | EPI1703041 | K | I | D | L | F | D |
| A/South Australia/36/2019 | SIAT1 | EPI1387334 | E4 | EPI1440496 | T | G | D | P | F | D |
| A/South Australia/39/2019 | SIAT1 | EPI1387337 | E4 | EPI1440498 | K | G | D | P | S | D |
| A/South Australia/4/2019 | SIAT1 | EPI1371913 | E4+E2 | EPI1698473 | K | D | D | L | S | D |
| A/Sydney/53/2019 | MDCK-SIAT1+SIAT1 | EPI1430420 | E3+E2 | EPI1588472 | K | V | D | L | F | D |
| A/Tasmania/503/2020 | SIAT1 | EPI1752480 | E5 | EPI1848147 | K | R | D | L | S | D |
| | | | EX | EPI1868371 | K | R | E | L | F | D |
| | | | EX | EPI1868373 | K | R | D | L | F | D |

Continued

| VIRUS NAME | NON-EGG-PASSAGED | | EGG-PASSAGED | | MUTATION IN EGG-PASSAGED | | | | | |
|---|---|---|---|---|---|---|---|---|---|---|
| | PASSAGE HISTORY | HA ACCESSION NO. | PASSAGE HISTORY | HA ACCESSION NO. | 160 | 186 | 190 | 194 | 219 | 225 |
| A/Vermont/11/2019 | Original | EPI1428265 | E2 | EPI1439232 | K | G | N | L | S | D |
| A/Vermont/14/2019 | Original | EPI1428013 | E2 | EPI1439224 | K | G | N | L | S | D |
| A/Vermont/25/2019 | Original | EPI1618933 | E3 | EPI1713735 | K | V | D | L | S | D |
| A/Victoria/223/2019 | SIAT1 | EPI1584570 | E3 | EPI1610406 | T | G | D | P | F | D |
| A/Victoria/703/2019 | SIAT1 | EPI1430441 | E2 | EPI1444936 | K | G | D | L | S | G |

For passage history, ‘C’: passaged in cells. ‘E’: passaged in eggs. The passage number is indicated as the suffix. ‘X’ indicates missing information. Egg-adaptive mutations are highlighted in yellow.
